# Fitness Landscapes Reveal Context-dependent Benefits of Oviposition Choice

**DOI:** 10.1101/2021.05.27.445916

**Authors:** Vrinda Ravi Kumar, Gaurav Agavekar, Deepa Agashe

## Abstract

Resource choice behaviour has enormous fitness consequences and can drive niche expansion. However, individual behavioural choices are often mediated by context, determined by past experience. Are such context-dependent behaviours adaptive? Using *Tribolium castaneum* (the red flour beetle), we demonstrate that context-dependent oviposition choice reflects distinct, context-specific local fitness peaks. Manipulating female egg allocation in a habitat containing optimal and suboptimal resource patches, we measured offspring fitness to generate fitness landscapes as a function of all possible oviposition behaviours (i.e., combinations of fecundity and resource preference). Females from different age and competition contexts exhibit distinct behaviours which optimize different fitness components that are linked in a tradeoff. With increasing age and prior exposure to competition, they produce few but fast-developing offspring that are advantageous under high resource competition. In contrast, young naïve females produce many slow-developing offspring, beneficial under weak competition. Systematically mapping complete context-dependent fitness landscapes is thus critical to infer behavioural optimality and offers predictive power in novel contexts.

**Preprint available at** - https://www.biorxiv.org/content/10.1101/2021.05.27.445916v1.full

**Citation** - Vrinda Ravi Kumar, Gaurav Agavekar, Deepa Agashe; bioRxiv 2021.05.27.445916; doi: https://doi.org/10.1101/2021.05.27.445916

## INTRODUCTION

Organisms often encounter heterogeneous habitats, and their behavioural choices in such situations ultimately determine their fitness. This is perhaps best exemplified by females’ reproductive decisions when faced with alternate resources. For example, females of phytophagous insects can distribute their eggs on different host plants, which affects offspring survival and growth and determines their reproductive success (1–2). Therefore, oviposition preference is predicted to face strong selection on par with other life history traits (3), sometimes leading to genetic correlations between preference and performance (2). However, oviposition choice can also vary dramatically with females’ prior experience, reflecting the ecological or internal context determined by their environment or life history (4). For instance, females of the polyphagous moth *Spodoptera littoralis* that develop on a specific host plant as larvae also prefer it subsequently for oviposition (5). More generally, ecological factors such as age and resource competition change across individuals’ lifespan, altering key components of oviposition behaviour such as fecundity (6–9) and oviposition choice (10–11). Given the potentially enormous range of contexts experienced by different individuals across generations, how do these context-dependent oviposition behaviours evolve?

Broadly, two kinds of selection pressures may drive context-dependent oviposition choice. First, female choice may evolve under selection for a single globally optimal (context-independent) preference, with context-dependent deviations from this optimum reflecting weak selection, environmental constraints or tradeoffs. For instance, limited perception of resource quality (12) or other search-related constraints (2, 13, 14) may lead to suboptimal preference, as could tradeoffs between maternal and offspring fitness (15) or the presence of competitors or predators (16–17). Alternatively, context-specific female choice may reflect context-specific selection to optimize distinct fitness peaks, with variable female choice perfectly attuned for each context (18). To our knowledge, no study has distinguished between these alternative hypotheses, perhaps because it is challenging to comprehensively connect past context to future fitness. For instance, to identify context-specific fitness optima it is critical to measure multiple fitness components, because many traits contribute to fitness and may trade off with each other (19–22) across contexts. Further, to correctly infer whether an observed behaviour is locally or globally adaptive (or maladaptive), we must generate a unified fitness landscape that encompasses all possible behavioural choices, including those outside the range of behaviours observed in nature. Using females from diverse contexts to create complete fitness landscapes also allows the prediction of female fitness in novel contexts. Illuminating the entire phenotypic fitness landscape is thus critical to understand the evolution of context-dependent oviposition behaviour.

We created such fitness landscapes for multiple fitness variables for the red flour beetle *Tribolium castaneum*, a widespread generalist pest of cereal flours that has long served as a model system in ecology and evolutionary biology (23, 24). Prior work on this species demonstrates significant context-dependent variation in key oviposition behaviours, modulated by age and population density (indicating strength of resource competition). Here, we explicitly connect prior female context, behavioural choice, and fitness, by conducting three sets of experiments focused on a choice between an optimal resource (wheat flour) and a suboptimal novel resource (finger millet flour) (Figure 1). We first validated the context-dependence of two aspects of oviposition behaviour (fecundity and preference) of females from six different age and density contexts (Experiment 1). Second, we generated complete fitness landscapes as a function of the full range of observed female fecundity and possible oviposition choices. To do this, we experimentally manipulated egg allocation in a habitat with both resource patches, and measured multiple components of offspring fitness (Experiment 2). Finally, we validated the fitness landscape by jointly measuring oviposition behaviours and consequent offspring fitness for females from three commonly-experienced contexts (Experiment 3). By placing oviposition behaviour in a comprehensive fitness landscape, we demonstrate that divergent, context-dependent oviposition behaviours could evolve via selection driving females towards context-specific local adaptive peaks for distinct fitness components.

**Figure 1.**
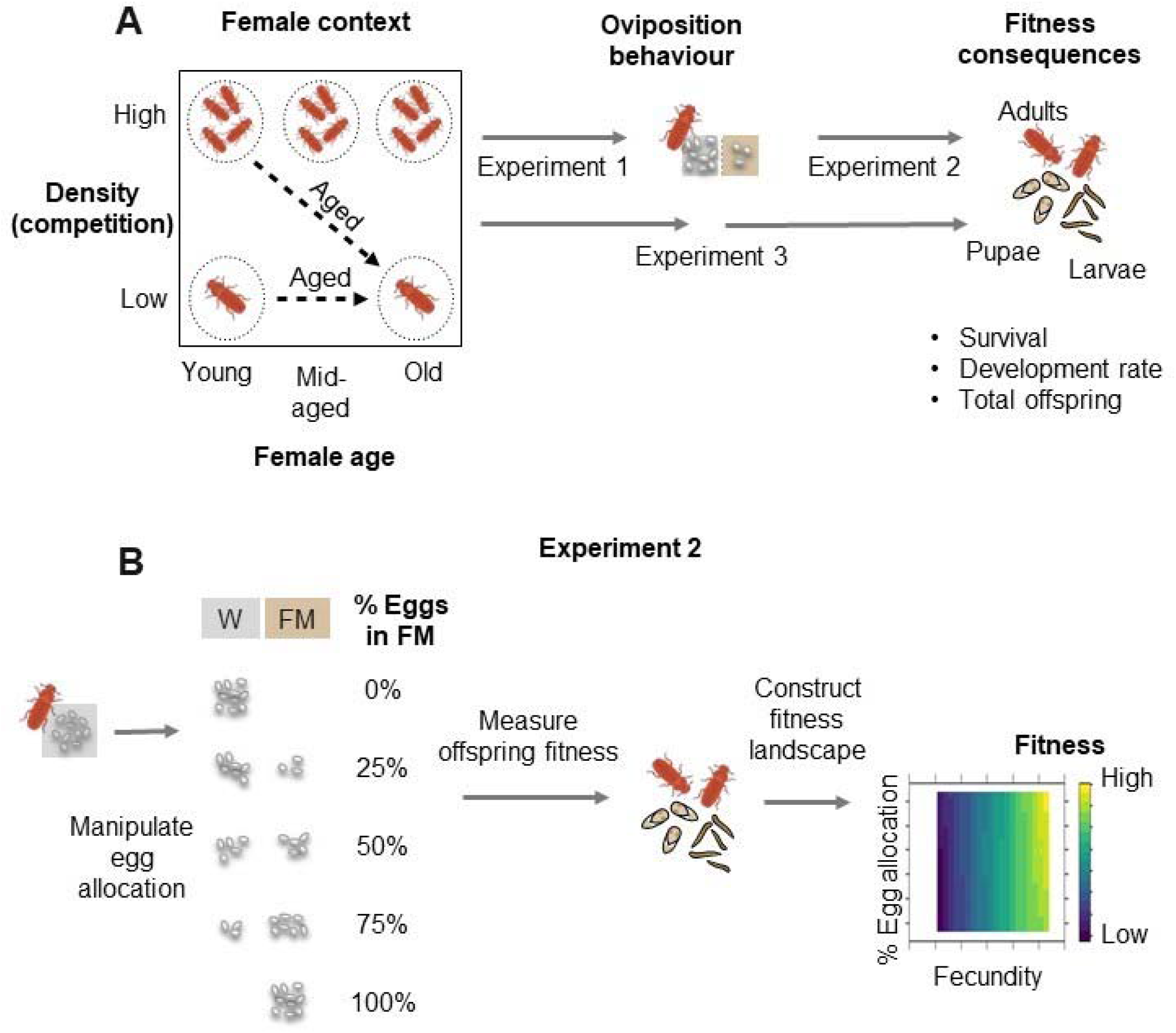
Experimental design. (A) Overview of key experiments (B) Details for experiment 2, used to generate fitness landscapes as a function of female fecundity and egg allocation across resource patches (W = wheat flour, FM = finger millet flour).

## METHODS

### Collecting females from different age and density contexts

We derived experimental individuals from previously described large, outbred laboratory “stock” populations of *T. castaneum* (25), provided with either wheat or finger millet flour. Flour beetles have highest fitness in wheat and show significantly slower development in finger millet (~37% slower development relative to wheat) (25). Details of population maintenance are given in SI methods. As required for different experiments (Figure 1), we generated females from six distinct age and density contexts (Figure S1). We used ‘young’, ‘mid-aged’ and ‘old’ females aged ~2 weeks, ~4 weeks and ~6 weeks after eclosion respectively (SI Methods), that experienced either low density (“LD”) or high density (“HD”) (Figure S1, SI Methods). Females from each context were mated with males from the same context to ensure consistency.

### Measuring oviposition behaviour and offspring fitness

To test how prior context changes oviposition behaviour, we measured the oviposition preference of females from different contexts (Figure 1, Experiment 1) for wheat vs. finger millet flour in individual 2-choice assays (SI Methods, Figure S2). We used the total number of eggs laid per female over 2 days to quantify fecundity, and the percentage of eggs deposited in finger millet as a proportional measure for oviposition preference for finger millet. The small oviposition window minimized egg cannibalism by larvae and adults. To measure the fitness consequences of female oviposition behaviours (Figure 1, Experiment 3), we removed adults and allowed offspring to develop undisturbed for 21 days without restricting larval movement, and quantified the resultant offspring fitness. We measured the proportion of eggs that survived three weeks of development (“survival”), the proportion of offspring that had successfully pupated or eclosed (“development rate”), and the total number of live offspring of all stages (“total offspring”). All measures of fitness were evaluated for the entire habitat, i.e., including both resource patches. Thus, we assessed the overall optimality of females’ reproductive strategies as they would operate in natural conditions, including all interactions between offspring and with their natal environment.

### Generating and validating fitness landscapes

To generate complete fitness landscapes (Figure 1, Experiment 2), we had to quantify the impact of the entire range of possible oviposition behaviours on offspring fitness. Hence, we collected females from three different contexts with distinct ranges of fecundity (young LD, young HD and old HD; see Results); but we did not allow them to express their oviposition choice. Instead, we collected eggs from each female and manipulated egg allocation systematically across wheat and finger millet patches in a 2-patch habitat (as in the oviposition choice assays), mimicking the full range of possible oviposition preferences (n=16-20 females per allocation). By distributing females with various fecundity values across egg allocation treatments, we dissociated female fecundity from resource preference, and then measured female fitness as described above.

Using these data, we generated fitness landscapes for each fitness component as a function of female fecundity (observed) and “preference” for finger millet (experimentally manipulated egg allocation). To do this, we extrapolated fitness values using local smoothing (‘loess’) by estimating the value of the three fitness measures for all possible combinations of female fecundity and oviposition preference using the ‘panel.2dsmoother’ function (26, from the latticeExtra package) in the levelplot function from the lattice package (27). Fitness was evaluated in an equally spaced 20 by 20 point grid across the sampled range of fecundity and preference. We also tested the impact of using a different smoothing function (generalized additive model, GAM) and altered grid size (ranging from 10 x 10 to 200 x 200 points).

These fitness landscapes allowed us to predict the consequences of any given combination of fecundity and oviposition preference. To validate the predictions (Figure 1, Experiment 3), we allowed females from three different contexts (young LD, young HD and old HD) to make oviposition choices; counted and returned eggs to the patch in which they were laid; and measured offspring fitness for each female. Using the median values of observed fecundity and preference for females from each context, we predicted their offspring fitness based on the fitness landscape and compared the predicted values with the median observed fitness values using a paired t-test.

### Statistical analysis

All analyses were conducted using R (28) in RStudio (29). We assessed the impact of age and density on female fecundity and of age, density and fecundity on oviposition preference (Experiment 1) and the impact of context, fecundity, and manipulated egg allocation on offspring fitness components (Experiment 2). We used ANOVA models for fecundity; generalised linear models with binomially distributed errors (‘binomial GLM’) for female oviposition preference, offspring survival and development rate; and a GLM with Poisson errors (‘Poisson GLM’) for the total number of offspring (SI methods).

## RESULTS

### Prior context alters female oviposition behaviour

Based on earlier work, we expected that female age and prior exposure to population density (an indicator of resource competition) would impact oviposition behaviour, i.e., female fecundity and preference for the novel finger millet resource (Figure 1, Experiment 1). Overall, fecundity ranged from 0 to 48 eggs, and oviposition choice ranged from 0 to 100% preference for finger millet. However, these traits differed significantly across females from different contexts (Figure 2A, 2B). For instance, female fecundity typically decreased with age and prior exposure to high population density (“HD”) (Figure 2A; ANOVA, F = 32.34, p < 0.001). Young females at low density (“LD”) had the highest fecundity, whereas HD females of the same age laid only half as many eggs (Tukey’s HSD, difference ~16 eggs, p < 0.001), similar to mid-aged and older females that experienced HD throughout their adult life (Tukey’s HSD, difference (mid-aged HD) = 15.5 eggs, p < 0.001; difference (old HD) = 18.5 eggs, p < 0.001). This broad effect of age was consistent when the same set of young females were allowed to age (Figure S4); and in an independent experimental block with young LD, young HD and old HD females (Figure S5). Concomitant with fecundity, oviposition preference for finger millet also typically decreased with age and density (Figure 2B), except for old HD females which oviposited randomly. Combining data across contexts, females with high fecundity laid more eggs in finger millet (Spearman’s rho = 0.303, p < 0.001; Figure 2C). However, fecundity and preference were not correlated within a given context (except for aged LD females; Figure S6 shows more experimental blocks).

**Figure 2.**
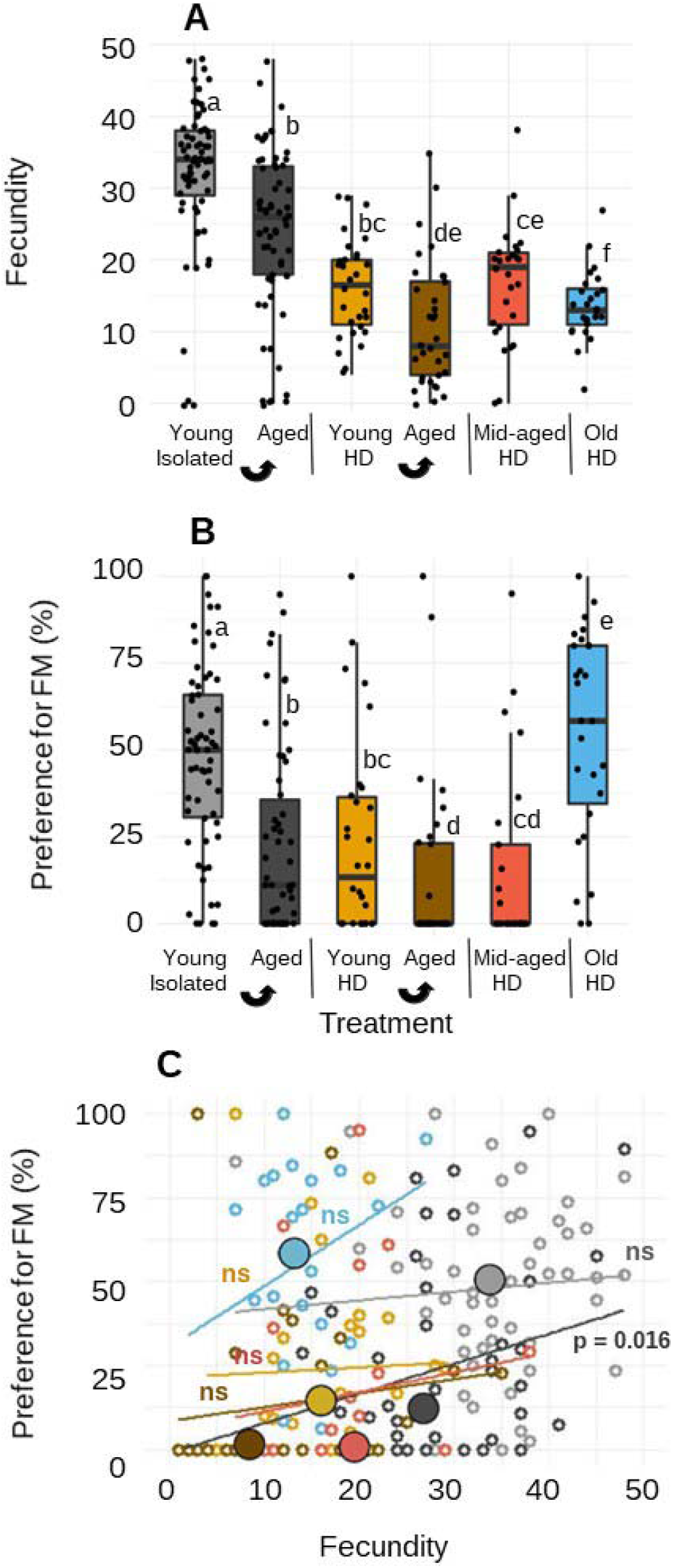
Oviposition behaviour of females from different prior contexts (Experiment 1). (A) Total fecundity and (B) % Eggs laid in finger millet (FM) in a two-patch habitat (LD: low density, HD: high density). Boxplots are coloured by female context; darker shades of a given colour represent the same set of females after ageing. Sample sizes are noted below boxes in panel A. Letters above boxes indicate results of a binomial generalized linear model, such that boxes with the same letter are not significantly different. (C) Correlation between total fecundity and preference for finger millet for females from each context (for aged LD females, Spearman’s rho = 0.315; ns = correlation not significant); lines indicate best-fit linear regressions for each context. Each point represents data for a single female; large filled circles indicate the median fecundity and preference for females from a given context; points and lines are coloured by context, as in panels A and B.

Despite these context-specific trends in oviposition behaviour, we noted some exceptions. When young HD females were aged, they continued to show a strong preference for wheat (typical of HD females) despite experiencing LD for 4 weeks (compare young HD vs. aged, Figure 2B), and also failed to exhibit the high fecundity commonly observed at low density (Figure 2A). Thus, the impact of early experience with HD seemed to last for a very long time. Additionally, old HD females did not prefer wheat as predicted by the age- and density- dependent trends in other females. Hence, preference may shift non-monotonically as females experience high density for different time intervals. Nonetheless, it is clear that age and density present appropriate contexts that influence two key components of oviposition behaviour. Therefore, we proceeded with measuring the fitness consequences of these divergent context-dependent resource preference behaviours.

### Fitness landscapes predict consequences of divergent oviposition behaviour

To quantify the fitness impacts of potential oviposition behaviours (Figure 1, Experiment 2), we allowed females to lay eggs in wheat and then manually distributed each female’s eggs in wheat and finger millet patches, covering the entire range of possible oviposition preference. To include the widest possible range of fecundity (Figure 2A), we sampled females from three different contexts (young LD, young HD, and old HD females), and measured three metrics of offspring fitness (Figure S7). This design ensured that we sampled the full range of possible fecundity– preference combinations to estimate their impact on fitness.

Across female contexts, offspring survival was largely invariable (~75% offspring survived). Interestingly, young LD females produced twice as many offspring as females from other contexts (~20 vs. ~10 per female), although they had the slowest development (~25% offspring completed pupation in 3 weeks, vs. ~50-75% for young HD and old HD females). Thus, distinct oviposition behaviours associated with different female contexts can dramatically alter offspring fitness. Notably, offspring development rate was strongly negatively correlated with female fecundity as well as total offspring (Figure 3A–B, Figure S8), reflecting a well-known density-regulated tradeoff between offspring number and development time in this species (23).

**Figure 3.**
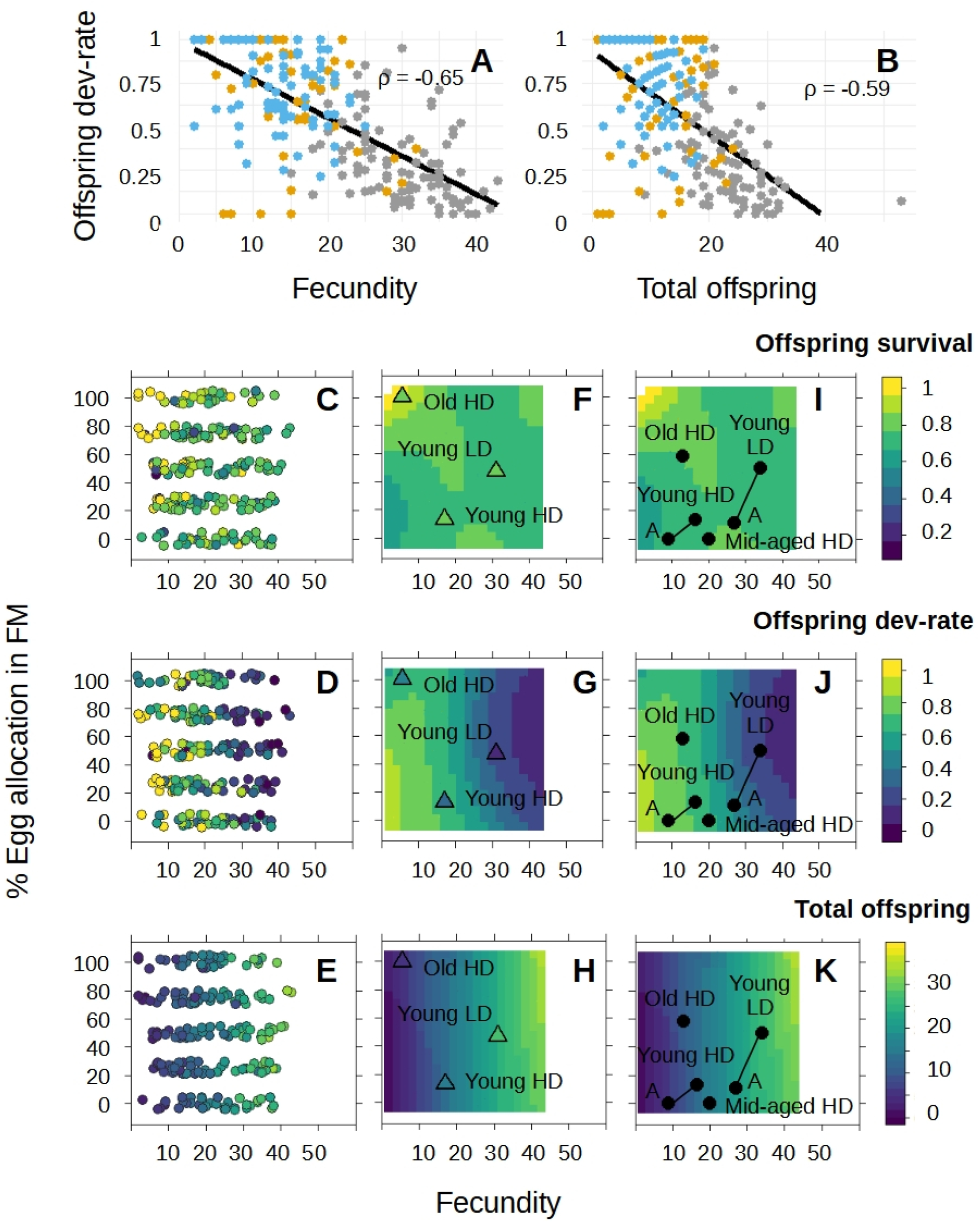
Complete fitness landscapes predict the fitness consequences of female oviposition behaviour in diverse contexts. Linear relationship between offspring development rate and (A) maternal fecundity and (B) total offspring (colour scheme as in Figure 2; n = 96 for young LD; n = 92 for young HD; n = 80 for old HD). Each point indicates data for a single female. (C-E) Measured fitness values (indicated by the z-axis colour bar on the right) for individual females for each egg allocation treatment (Experiment 2; n = 16-20 per egg allocation) spanning their natural fecundity range. Points are slightly jittered along the y axis for visualization. (F–H) Fitness landscapes generated for each fitness component using loess smoothing on data shown in (C-E). For validation, landscapes are overlaid with data from an independent experiment (Experiment 3) where we measured both oviposition preference (which determines the position of each triangle in the landscape) and offspring fitness (which determines the colour fill of each triangle). A close match in the colour of the triangle and its background indicates high predictability of fitness consequences (also see Table S1). (I–K) The same fitness landscapes overlaid with median observed oviposition behaviours for females from different contexts (obtained from Experiment 1, Figure 2), indicated by black filled circles. Lines connect cases where the same set of females were aged (indicated by “A”). The position of each circle shows the predicted fitness values for females from a given context.

To visualize the impacts of oviposition behaviour on fitness, we used data for each fitness component (Figures 3C-E) to construct complete fitness landscapes as a function of the two key aspects of oviposition behaviours (female fecundity and oviposition choice; Figures 3F-H). Since the landscapes encompass the entire range of possible oviposition behaviours (all possible preferences and the entire natural fecundity range), they offer critical insight into the fitness consequences of oviposition decisions for all females when offered a similar choice. Recall that females from different contexts exhibit distinct combinations of fecundity and preference (Figure 2), such that they occupy different positions in these landscapes.

As expected, the fitness landscape for survival was relatively flat (focus on the contours, ignoring triangles; Figure 3F). However, landscapes for the other two fitness components paint a more interesting dichotomy. Overall, most of the variation in fitness was distributed along the fecundity axis, suggesting (as expected) that fecundity is a powerful modulator of fitness outcomes. For development rate, the landscape unsurprisingly predicts a fitness optimum at low fecundity and low preference for finger millet; with rapid development at low fecundity regardless of preference (Figure 3G) suggesting that competition among offspring strongly impacts development rate. In contrast, offspring production is maximized at high fecundity (as expected) and strong preference for finger millet, with a relatively weak impact of preference (Figure 3H). These contrasting predicted fitness optima for offspring development rate and total number of offspring largely reflect the underlying negative tradeoff between the two components (Figure 3B, independent experimental blocks in Figure S8), primarily mediated by fecundity (Figure 3A). The estimated fitness maxima predicted by the landscapes were robust to changes in the fitting method (loess vs. GAM) and the grid density over which fitness was evaluated.

Separate fitness landscapes constructed for each female context generated similar patterns, indicating that the unified fitness landscape is robust (Figures S9, S10). We also confirmed that the landscapes accurately predict the fitness consequences of any combination of female fecundity and oviposition choice. In an independent experiment, we allowed females to express their oviposition preference (used to predict fitness) and measured the fitness consequences of their choice (Figure 1, Experiment 3). Observed female fitness matched predictions from the original landscape (compare the triangle colour with background colour, Figures 3G–I; Table S1, p > 0.05 for all three fitness parameters), even though the oviposition behaviour of old HD females in this experiment was different from what we had observed in Experiment 1 (comparison shown in Figure S5). Thus, fitness landscapes generated by scrambling the inherent association between female fecundity and preference and fitness accurately predicted the fitness consequences of diverse oviposition behaviours.

### Females from different contexts use alternate adaptive oviposition strategies

The fitness landscapes generated above suggest two general oviposition strategies that each increase different fitness components – high preference for wheat with low fecundity, or high acceptance of finger millet with high fecundity. To explicitly connect prior female context with these strategies, we asked whether context-dependent behaviours are consistent with either of these predicted fitness peaks. To do this, we used the observed context-specific oviposition behaviours described in Figure 2, which represent females from diverse age and density contexts. We predicted the fitness outcomes for each context by overlaying the median values of female fecundity and preference onto the landscapes for each fitness component. As expected, females from different contexts were positioned widely across the landscape (Figures 3I-K). As noted earlier (Figure 2A), females that were older or experienced high density generally had low fecundity; whereas younger females or those at low density laid many more eggs. At the same time, they also had divergent oviposition choices. In light of the fitness landscapes, we can connect these contrasting patterns, now revealed as alternate adaptive strategies. For instance, young LD females were close to the peak for total offspring, whereas young HD females produce fewer, rapidly-developing offspring. Distinct oviposition behaviours thus place females close to different locally adaptive fitness peaks.

Although the fitness landscapes were generated using females from only three contexts, we could also use them to predict fitness for other contexts, using only their oviposition behaviours. Strikingly, females from the two “aged” contexts also present divergent outcomes, though both (young HD and young LD females) were aged for 4 weeks under identical LD conditions. Their distinct positions in the fitness landscapes are driven primarily by their distinct fecundity, indicating a long-lasting impact of early exposure to high density. Aged young HD females showed a dramatic reduction in offspring production (Figure 3K), but their increased preference for wheat increased offspring development rate (Figure 3J). In summary, the complete fitness landscapes reveal that maternal context alters distinct fitness components by differentially influencing two key aspects of oviposition behaviour.

## DISCUSSION

Here, we present the first comprehensive analysis of the linkages between females’ prior context, their oviposition behaviour, and offspring fitness. By governing female behaviour, the contexts determine females’ placement on the fitness landscapes, defining their fitness. Thus, a context-specific interpretation of fitness is critical to infer optimality of specific behaviours. Importantly, to identify distinct local fitness peaks we need separate landscapes for each fitness component. For instance, if we only measured offspring development rate, we would incorrectly conclude that young LD females behave maladaptively. However, using the ecological selection faced by females in their prior context (here, high vs. low competition) we can better assess the adaptive consequences of their behaviour. We thus infer that under conditions of resource abundance (when their offspring are unlikely to face strong competition), females can effectively maximize the total number of offspring even if they develop slowly. Such context-dependent optimization of different fitness components may explain prior cases of apparent maladaptive resource use. For instance, flour beetles exhibited seemingly maladaptive density-dependent niche contraction in a heterogeneous habitat containing wheat and corn (a poor resource) (31). Based on our current results, we suggest that the observed preference for wheat at higher densities may reflect optimization of offspring development rate or survival; whereas at low density, females maximize total offspring production. Thus, age and density contexts primarily alter female fecundity, and females accordingly modulate their oviposition resource choice to optimize either total offspring or offspring development rate. Thus, our results demonstrate the lack of a single global fitness peak for oviposition behaviour, and show that a change in context can shift female behaviour towards alternative fitness peaks.

How generalisable is the suggested coupling between context-specific behaviour and fitness? Here, we considered two very broadly applicable contexts across species – age and experienced population density (competition) – that differ both between individuals in a population and across an individuals’ lifespan. In *Tribolium*, transitions across contexts are expected to occur in association with population growth and dispersal events (32–35). Ageing is universal and change in the strength of competition is a common feature of the life history of many organisms; both factors can impact oviposition. Mechanistically, any change in context that alters female body condition could influence resource allocation towards competing aspects of fitness (reflecting tradeoffs), ultimately determining oviposition behaviour (36–38). Hence, we can broadly frame our results to make generalized predictions about the context-dependent optimization of fitness for any species (Figure 4). While the exact fitness components that face selection or trade off across contexts may vary across species or environments, we suggest that the core axes that determine the impact of context on behaviour – age and resource competition – should be widely applicable.

**Figure 4.**
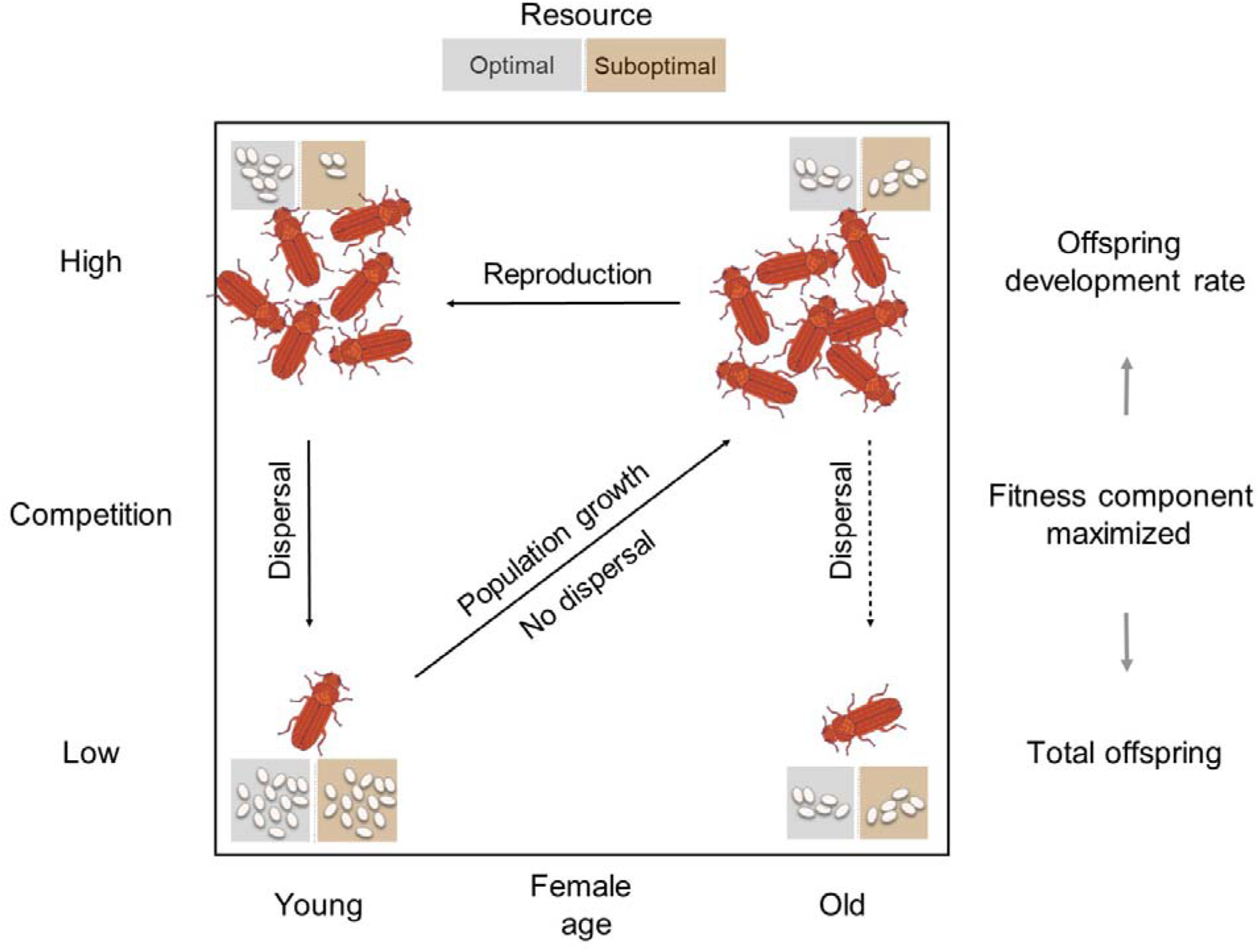
Illustration of the proposed general linkages between female context, oviposition behaviour and fitness. Young females that experience low competition (as a function of low population density; bottom left) may reproduce and establish a population as they age, increasing population density and competition (top right). Further reproduction will generate young females that also experience high competition (top left). Density-dependent dispersal may then expose females to low competition while colonizing a new habitat (bottom left). Old females may also disperse away from high density conditions (bottom right), though we do not have data on this aspect (indicated by dashed line).

Our results show that females in different contexts can optimize different fitness components that trade off with each other. How does preference evolve, given that females’ behaviour is pulled in opposing directions across their lifetime? Broadly, frequent changes in selective context during the evolutionary history of a species could lead to the observed strong, specific associations between female context and expressed oviposition behaviour. We speculate that in *T. castaneum*, instead of selection acting independently on both fitness components, only offspring development rate evolves under strong selection. This is generally expected for species with a high intrinsic growth rate that often experience high population density, such as pests and invasives (39,40). At different times during their life (depending on age and density), female beetles may thus experience either intensified or relaxed selection to produce fast-developing offspring. For instance, prior work shows that larval development time in *Tribolium* increases under high larval or adult density (41,42). Thus, one way to increase offspring development rate under high resource competition is to decrease fecundity, and *T. castaneum* females may employ this strategy (43). Such selection on development rate may additionally drive high preference for optimal resources (here, wheat) under high competition. But how do we explain the random resource preferences of both old HD females and young LD females in this framework? One possibility is that old females – especially after prolonged exposure to high density – are unable to make effective choices, and therefore allocate eggs randomly. Meanwhile, under low competition (e.g., young LD females), selection on offspring development rate should be weak, allowing females to randomly allocate their eggs across habitats. Thus, distinct mechanisms may explain the convergent random preference behaviour of young LD and old HD females; this hypothesis needs to be explicitly tested. Nonetheless, our data suggest that both age and density have immediate impacts on fecundity, in turn altering female choice. Hence, fecundity and oviposition choice are inextricable, whereas most prior work on female context has focused on fecundity alone (38,43,44). We suggest that for dispersing species that often encounter heterogeneous habitats, joint measurement of fecundity and resource choice is essential to fully understand oviposition behaviour.

Our results also suggest a long-lasting impact of prior context on behaviour. Aged young HD females – despite a 4-week exposure to low density – continued to produce fast-developing offspring (optimal at high density), instead of increasing fecundity to maximize total offspring (optimal at low density). The failure to increase fecundity in response to a relaxation in adult competition is costly (~15 fewer offspring than expected under low density) and surprising (45), because flour beetles routinely experience changes in population density that drive their population dynamics (46) and dispersal (32,33,35). As a result, we expect flour beetles to experience strong selection to rapidly respond to changes in competition and alter oviposition behaviours. We speculate that early physiological changes due to high density (47–51) may be irreversible, preventing females from optimizing their oviposition behaviours even under favourable conditions. Such “carry-over effects” are widespread in the animal kingdom (52–54). Alternatively, the suboptimal suppression of fecundity may persist in the population due to weak selection, e.g. if old isolated (i.e., recently dispersed) females are rare in the normal life cycle of *T. castaneum* (33). Further work is needed to understand this interesting phenomenon and distinguish between the various hypotheses.

Context-dependent oviposition behaviour has longer-term implications for species’ evolution. For instance, reduced female preference for the ancestral or optimal resource could initiate the process of niche expansion by exposing offspring to selection in a new habitat, or by influencing the resource preference and performance of subsequent generations. We showed previously that in flour beetles, larval experience with novel resources increases later use of the same resources (25); a pattern that is also observed in many other insects (55–58). Hence, initial female acceptance of a new resource can increase its use by offspring, exposing offspring to selection and potentially improving their performance over time. Therefore, preference can lead and performance may follow when species colonize new habitats (59–61). More generally, random or no-preference behaviours have been suggested to drive the evolution of generalist species (62–64). Our work indicates that such effects may occur only when females from specific contexts colonize new habitats (in our case, old HD and young LD females). Hence, the prior experience and history of colonizing females may be important determinants of successful niche expansion.

In conclusion, we suggest that it is important to quantify the linkages between female experience, body condition, oviposition behaviour, and offspring fitness to enable context-informed interpretations of optimality. Understanding these links is important to successfully use oviposition behaviours for pest management, such as in the push-pull strategy (65), where experience-dependent behaviour is exploited to reduce oviposition on crops. Although a large body of prior work has partially addressed these links, we have lacked predictive power in new environmental or individual contexts. Here, we show an effective and reliable way to generate broad predictions about the fitness consequences of female oviposition behaviours. By dissociating females’ preference from their fecundity, one can systematically cover the entire range of phenotype values that comprise the fitness landscape, including combinations of fecundity and preference that may not be directly or easily observed. One can then predict the fitness consequences of female behaviour in new contexts, without the need to measure fitness in each scenario. Studies documenting evolutionary traps already highlight the power of explicitly considering longer-term historical context to understand behaviour (66,67). We suggest that considering short-term female context is similarly important to understand female behaviours as drivers of niche expansion.

## ACKNOWLEDGEMENTS

We thank Laasya Samhita, Pratibha Sanjenbam, Rittik Deb, Shyamsunder Buddh, Sneha Garge and other members of the Agashe lab for discussion and comments on the manuscript. We thank Sumita Nanda for the beetle illustration used in figures. We acknowledge funding and support from the National Centre for Biological Sciences (NCBS-TIFR), the Department of Atomic Energy, Government of India (Project Identification No. RTI 4006) and a SERB Women Research Excellence Award (WEA/2020/000030) to DA.

