## Supplementary Information for "Fitness Landscapes Reveal Context-dependent Benefits of Oviposition Choice"

**SUPPLEMENTARY METHODS**

**Population maintenance**

We held all populations and experimental individuals at 33±2°C in a dark incubator, except during experimental handling under white light at room temperature (never exceeding 1 hour). To remove any prior insect infestation from the resources used for all maintenance and experimental assays, we cold-sterilized flour for at least 4 hours at -80°C and allowed it to return to room temperature before use.

**Collecting females from different age and density contexts**

*Age contexts*

Females become sexually mature within one week of eclosion and lay eggs continuously for at least 40 weeks (23, 30). However, fertility begins to decline starkly ~7 weeks post eclosion (29). Hence, we used 'young', ‘mid-aged’ and ‘old’ females aged ~2 weeks, ~4 weeks and ~6 weeks post-eclosion respectively, corresponding to a total of approximately 27% decline in fertility (measured for isolated, singly-mated females; 9).

*Density contexts*

For the low-density contexts, we held females in isolation (if they had mated previously) or with a single male (if the females had not mated previously), whereas high-density (“HD”) females were taken directly from stock populations (and had likely multiply mated). For the two “aged” contexts, after the first oviposition assay (described below) we allowed young LD and young HD females to age for an additional 4 weeks in LD conditions.

*Individual treatments*

1. Young low-density (“LD”): from the stock population, we removed pupae and identified their sex. We isolated each pupa in a 1.5 mL Eppendorf microcentrifuge tube containing ~1 g wheat flour for 11 days, and then paired adults for mating in a fresh tube with ~1 g wheat flour for. After 4 days, we immobilized individuals with a cold shock, sexed them under a microscope, and allowed females to recover in the incubator in fresh flour for 3 hours before using in experiments.

2. Young high-density (“HD”): we set up separate stock populations, and instead of propagating them, we allowed newly eclosed adults to continue to mate and oviposit in the same box for ~2 weeks. We then sexed and removed females and allowed them to recover in the incubator, before using in experiments.

3. Mid-aged HD: as above, ~4 weeks after adult eclosion.

4. Old HD: as above, ~6 weeks after adult eclosion.

5. Aged young LD: after the first oviposition assay, we paired young LD females with the same male as earlier in fresh flour, and allowed them to age for ~4 weeks (i.e., for the same time as the “old” females). We replaced the tube and flour every 3-4 days to minimize crowding due to offspring.

6. Aged young HD: after the first oviposition assay, we isolated young HD females in fresh flour and allowed them to age as above.

**Oviposition assays**

The assay habitat consisted of petri-plates (60 mm diameter) sawed in half. We fastened these two halves together with transparent tape to ensure a seamless fit. To ensure females encountered a uniform texture while moving around the plate, we placed a well-fit paper cutout over the base of the assembled petri-plate. We placed ~1g of each resource (double-sifted) at diametrically opposite locations in the petri-plate, ensuring that the resource piles were not in contact. To start the assay, we released a female in the center of the petri-plate, facing away from both resource patches (Fig S2), with finger millet randomly occurring on either side across replicates. Then we covered the petri-plates and placed them in the incubator for 43-45 hours. We removed the female and counted the number of eggs in each resource patch. We either discarded or replaced the females in the incubator in fresh tubes to age further (for the “aged” groups).

*T. castaneum* eggs hatch in ~48 hours; therefore, all eggs from all replicates were counted within 3-5 hours of removing the female. To facilitate egg collection, we pre-sifted flour twice through a fine mesh sieve (pore size 50 μm, Daigger Scientific USA) to remove large flour particles. To count the number of eggs in each resource patch within a single plate, we cut the paper base using a paper cutter. After this, we emptied the resource in each half-plate over a fine-mesh sieve to separate the eggs. We counted the number of eggs in each resource patch per female. We used the total number of eggs in the habitat (i.e., across both resource patches) to quantify fecundity, and the percentage of eggs deposited in the finger millet patch as a measure for oviposition preference for finger millet. If the flour patches were mixed during handling such that we could not reliably assign each egg to a single flour patch, we discarded the replicate.

In addition, we tested whether females explored both patches in this assay and oviposited randomly, as expected when both the patches comprised of the same resource. Individual females showed random oviposition preference in 2 independent blocks (Figure S3). Within each block, individual oviposition preference was random in homogenous habitats containing two patches of either wheat (fraction of eggs in a focal patch: Block 1, n = 15, 0.41-0.65, 95% CI of preference estimated by a binomial GLM; Block 2, n = 11, 0.47-0.55, 95% CI) or finger millet (Block 1, n = 19, 0.48-0.54, 95% CI; Block 2, n = 17, 0.44-0.54, 95% CI). Additionally, individual preferences of the tested females were normally distributed (Shapiro-Wilk test, p>0.05 for all distributions). Therefore, females typically explored and oviposited in both patches in this habitat.

**SUPPLEMENTARY FIGURES**

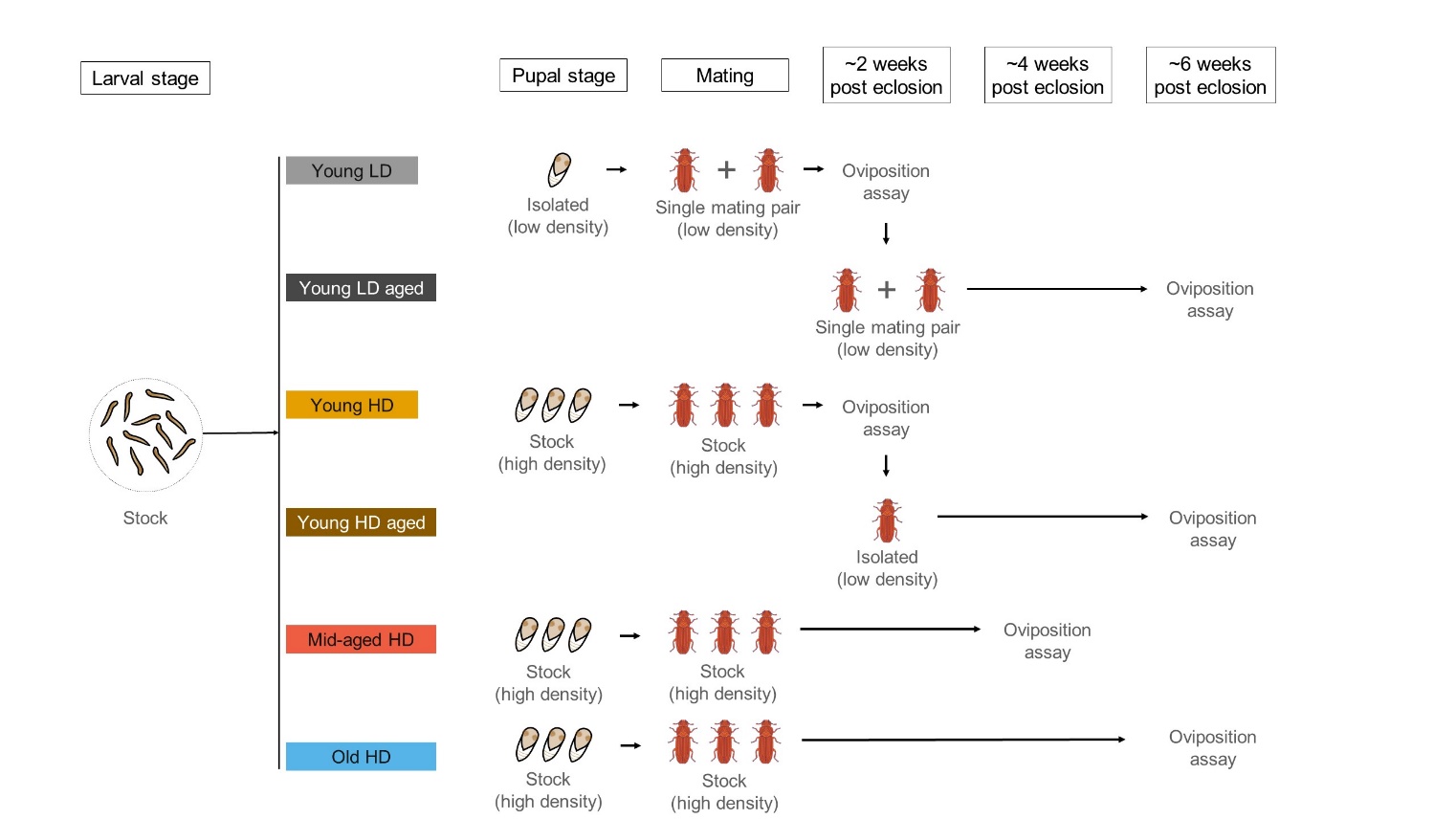

**Figure S1.** Schematic representing the different female contexts tested in Experiment 1 as a function of the different density treatments administered to them as they aged.

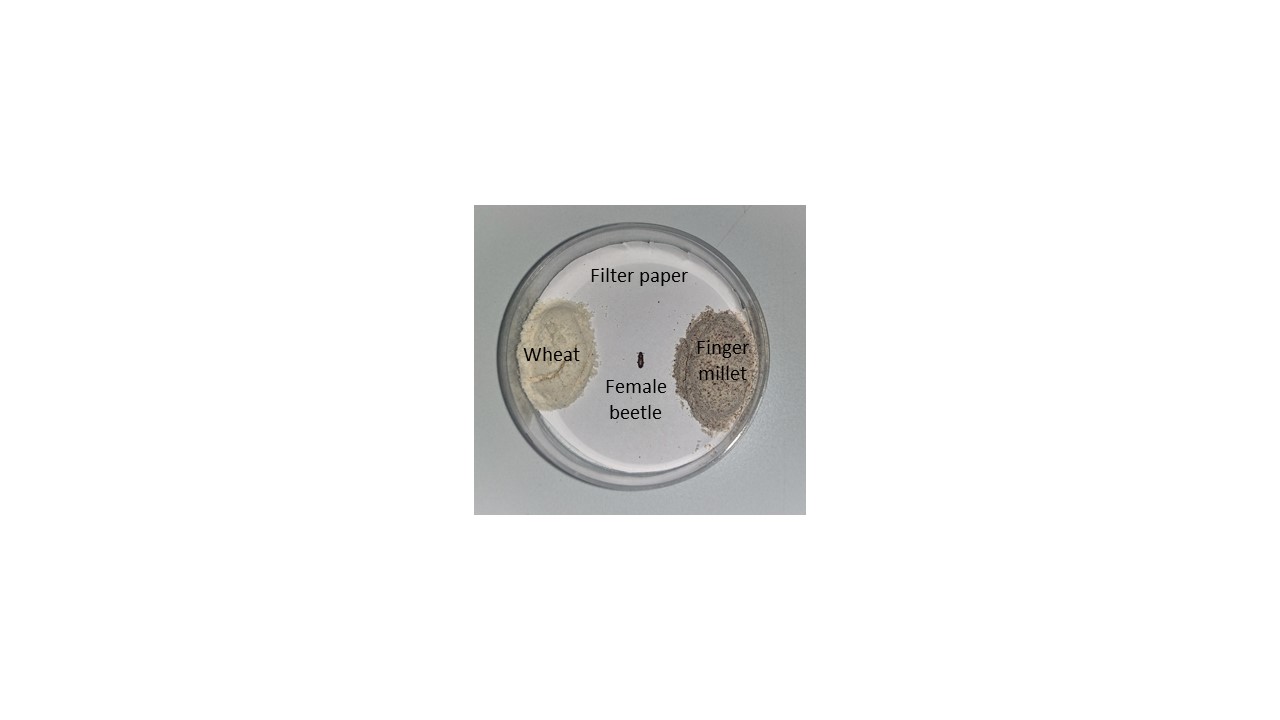

**Figure S2.** Picture of an oviposition assay plate, with an adult female placed in the center at the start of the assay.

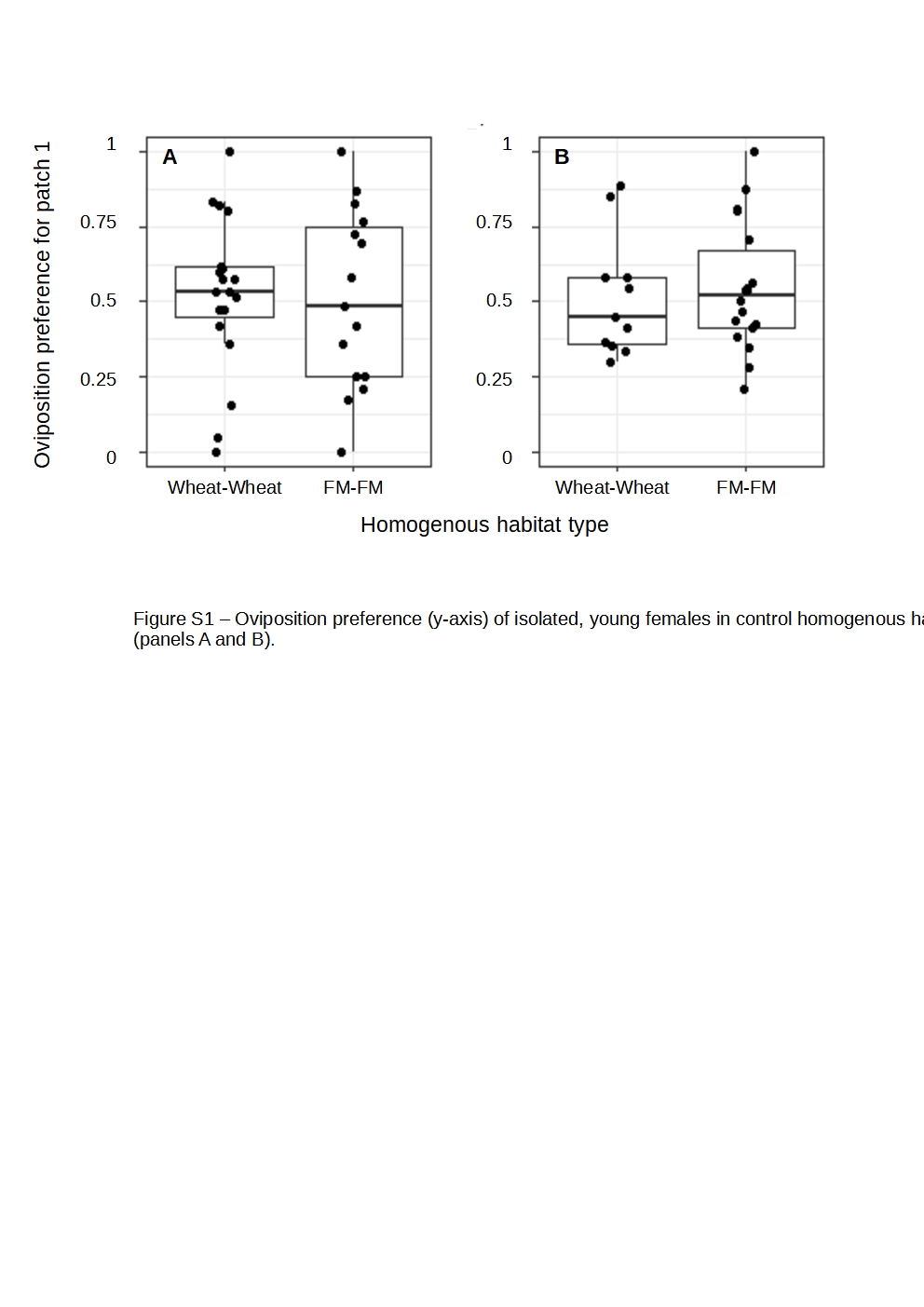

**Figure S3.** Oviposition preference (y-axis) of young LD females in control homogenous habitats where both patches were either wheat (n = 19 & 11 respectively) or finger millet ("FM"; n = 15 & 18 respectively) (x-axis). Preference was measured for 'patch 1' which was pre-marked on the plate before the start of the assay. These control females were tested in 2 independent experimental blocks (panels A and B). Relevant details are in SI methods.

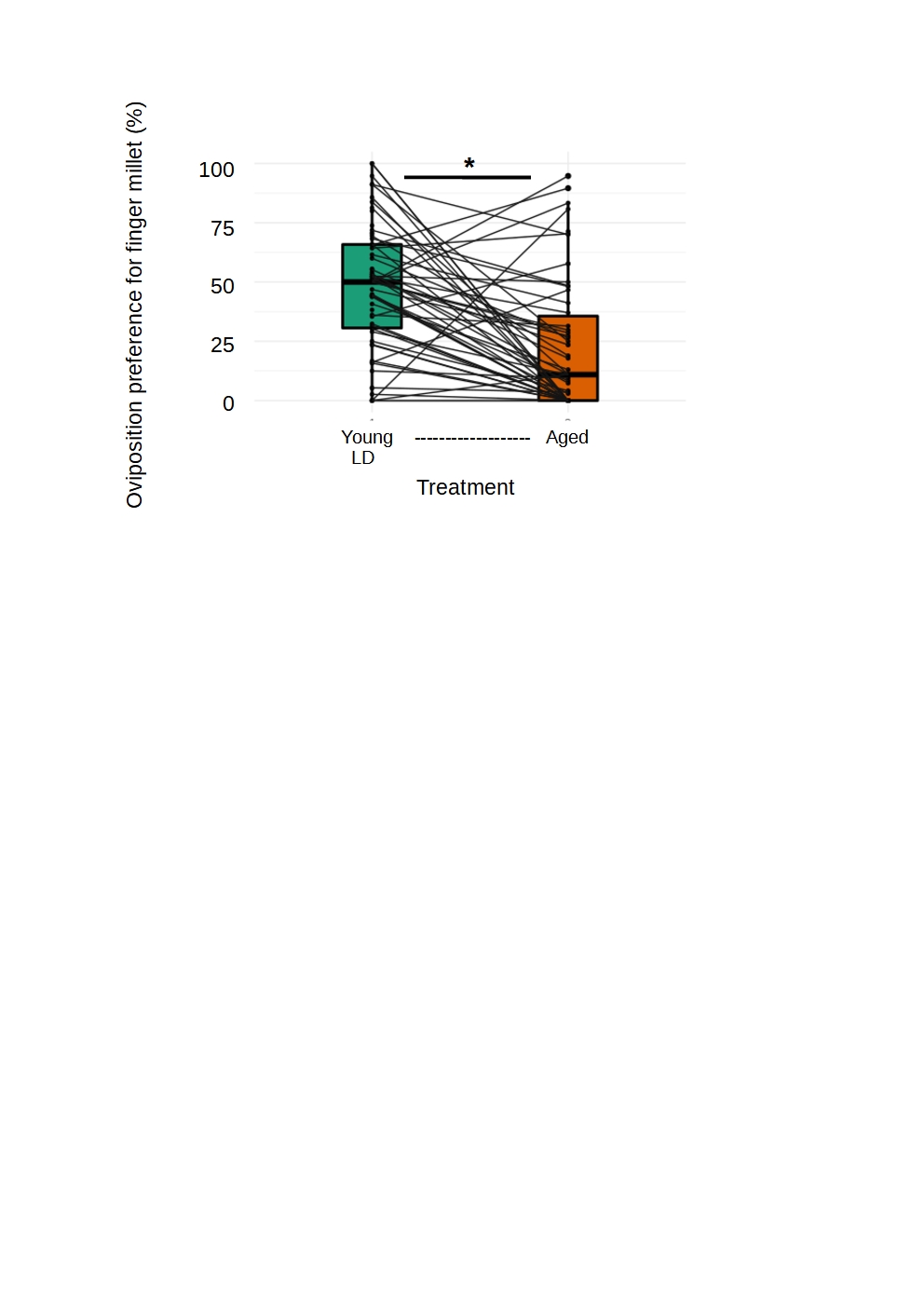

**Figure S4.** Impact of age on female oviposition choice. Young LD females (shown in Figure 2) were aged with a single male for ~4 weeks and then re-tested for preference. Lines connect the preference of the same female. A paired t-test shows a significant impact of ageing (mean of differences = 0.25, p < 0.001). n = 62 (young LD) and 57 (post-ageing) respectively.

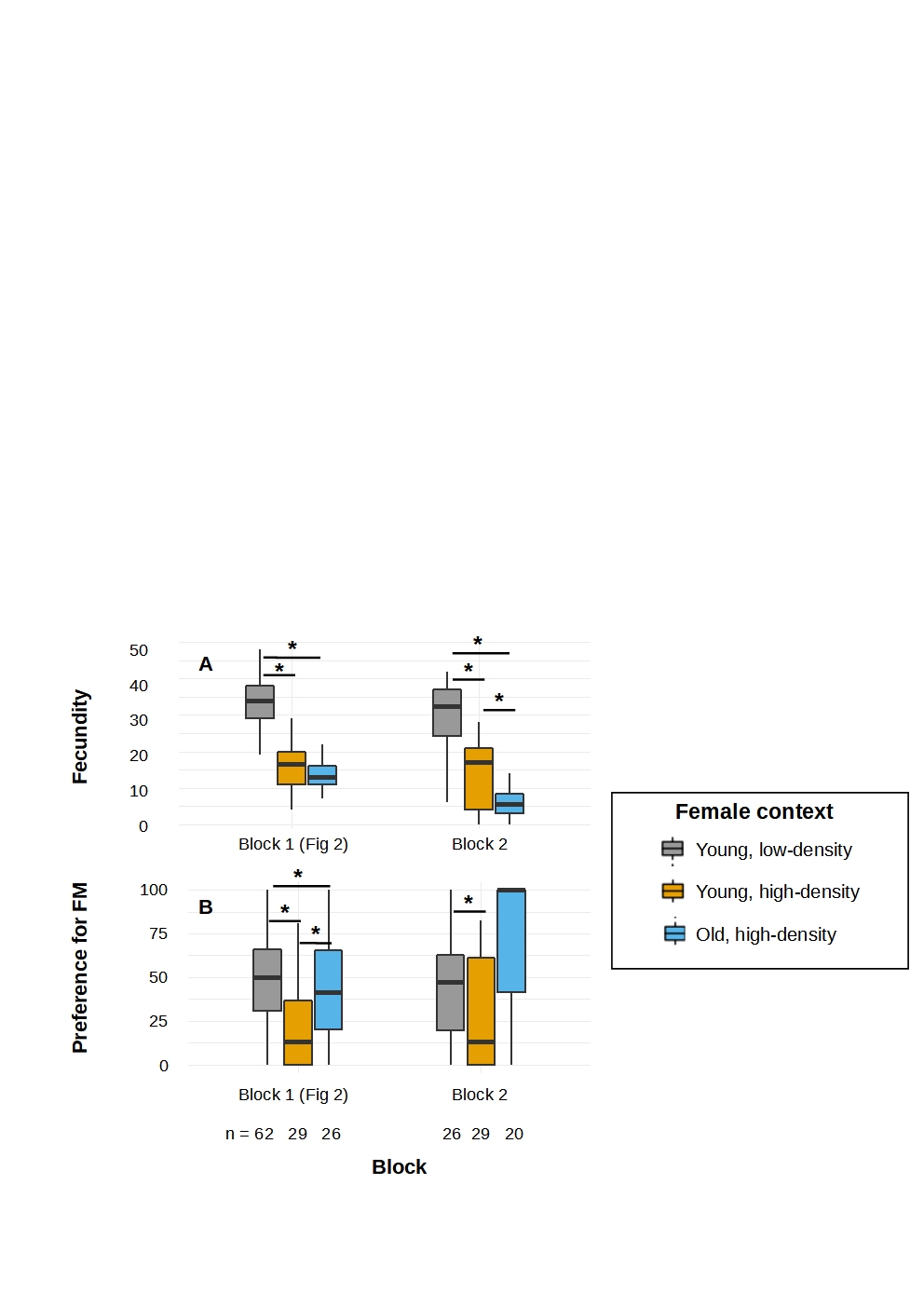

**Figure S5.** (A) Fecundity and (B) oviposition preference for finger millet (“FM”) of young LD (grey), young HD (orange) or old HD (blue) females in two independent experimental blocks (first block: Experiment 1; second block: Experiment 3). Sample sizes are indicated in panel B. These data were not combined with the data shown in Figure 2.

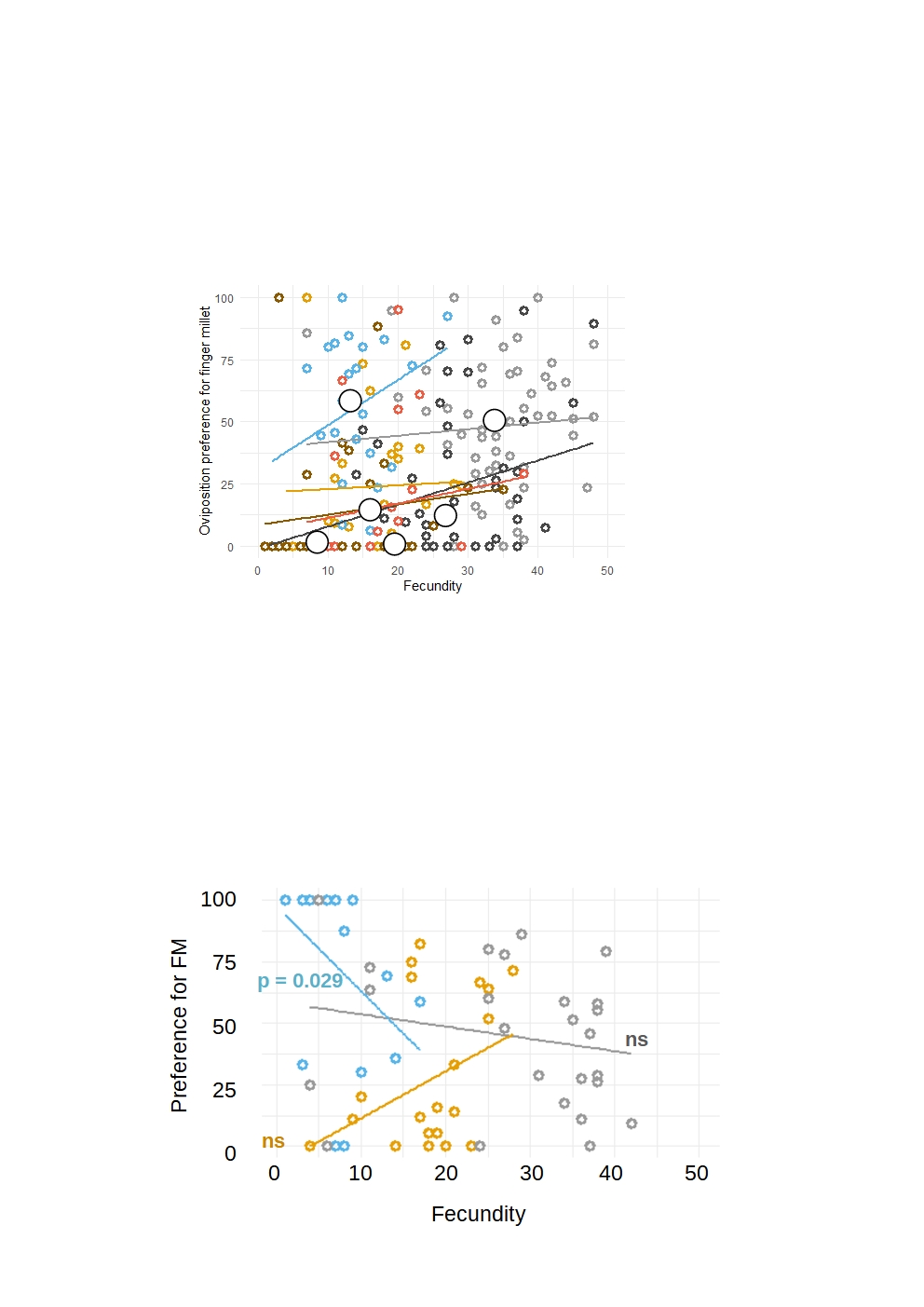

**Figure S6.** Correlation between total fecundity and preference for finger millet for each female group (Spearman’s rho for old HD females = – 0.513; ns = correlation not significant) from the second experimental block (Experiment 3). Colour scheme as in Figure 2; grey = young LD (n = 24), yellow = young HD (n = 20) and blue = old HD females (n = 14). Lines indicate best-fit linear regressions for each group.

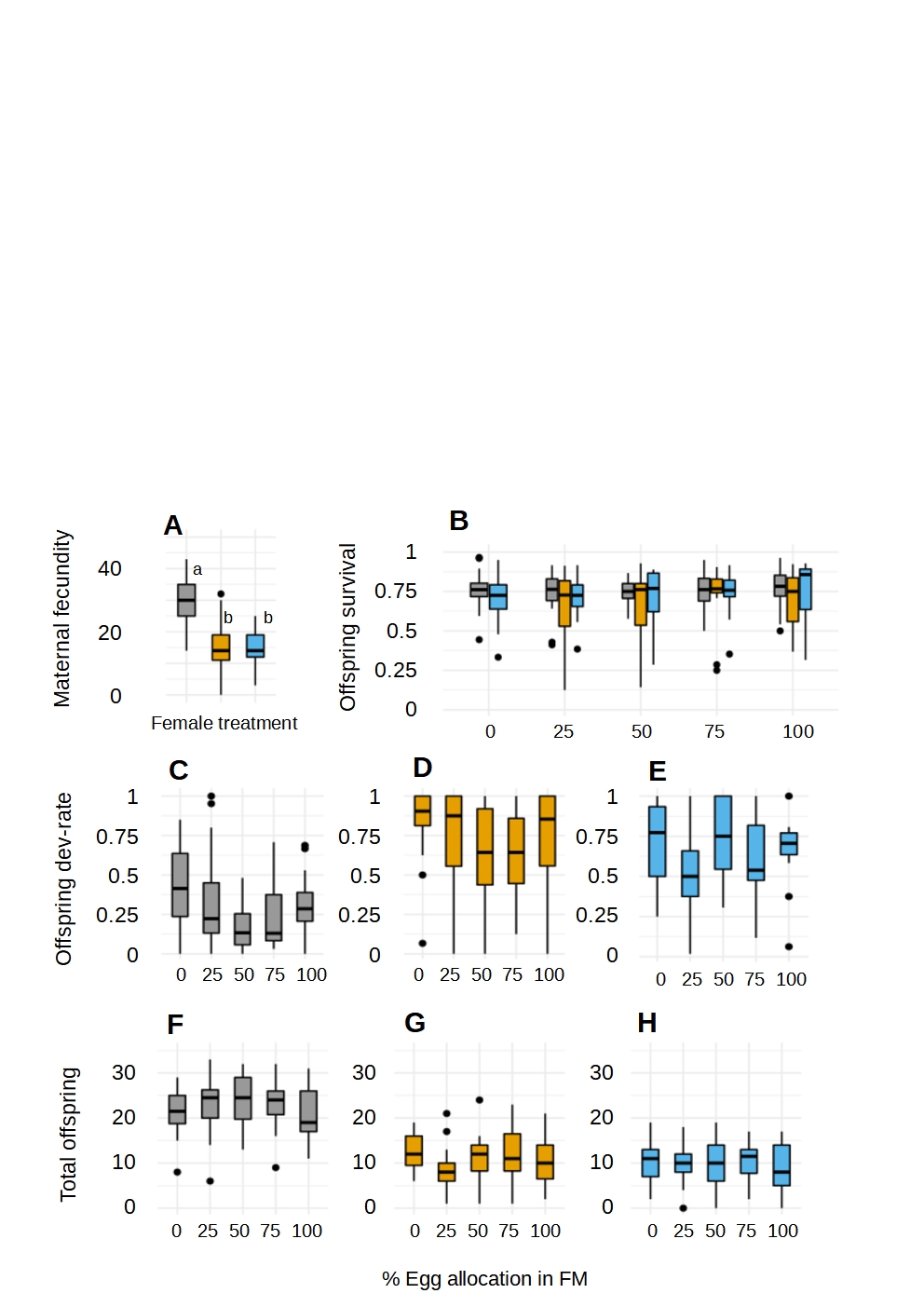

**Figure S7.** **Offspring fitness as a function of (manipulated) egg allocation across resource patches (Experiment 2).** (A) Female fecundity varied with female context. Boxplots marked with the same alphabet are not significantly different from each other as estimated by a Tukey HSD. Offspring fitness (n = 16-20 per egg allocation), estimated as (B) proportion of eggs that survived (C-E) proportion of surviving offspring that had completed development to pupation or adulthood and (F-H) total number of adult offspring. Boxplots are coloured by different female contexts, as in the key in Figure 2.

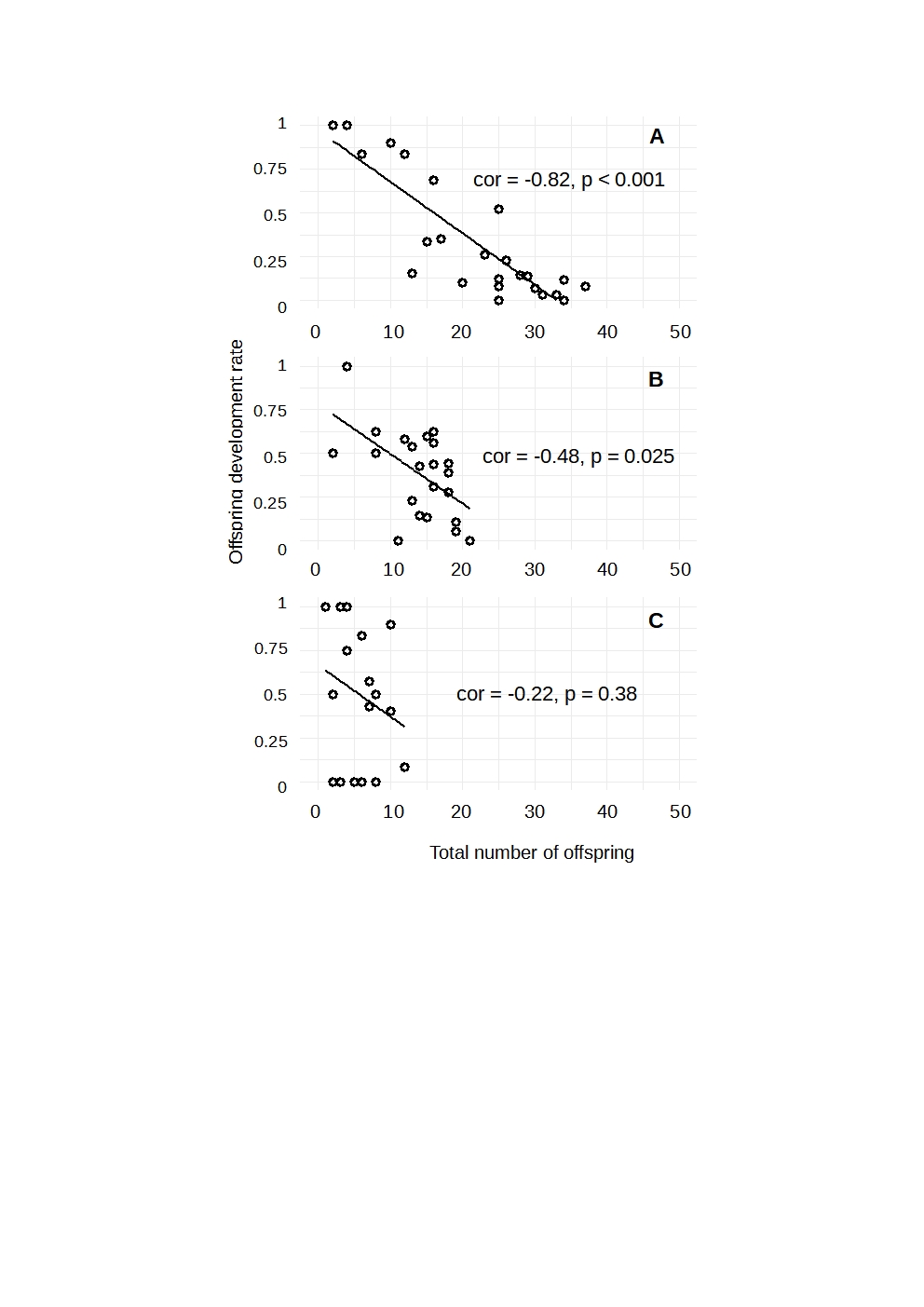

**Figure S8.** Linear relationship between offspring development rate and total offspring for (A) young LD (n = 24), (B) young HD (n = 22) and (C) old HD (n = 17) females. Each point represents individual females (from independent replication experiments) in the heterogenous wheat-FM habitat. The black line represents the linear regression between the two variables.

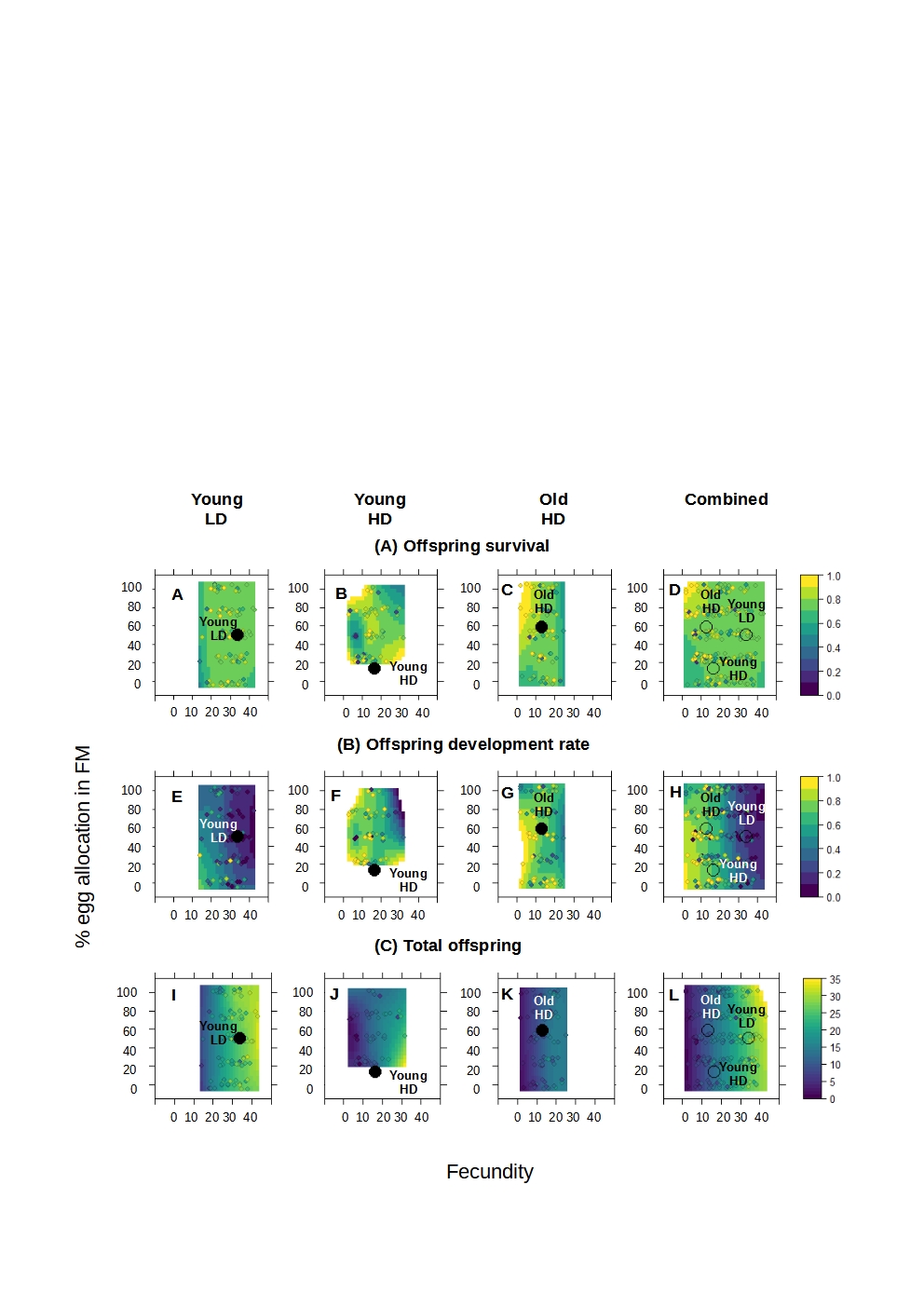

**Figure S9.** Estimated fitness landscapes for (A-D) offspring survival, (E-H) offspring development rate and (I-L) total number of offspring for individual female treatments shown in Figure 2. Fitness landscapes estimated individually for (A, E, I) young, isolated females (B, F, J) young females, (C, G, K) old females that developed under high density or (D, H, L) all combined. Contours were estimated using the loess function. Overlaid points represent the median preference-fecundity value for females in the respective treatment. In panels (D, H, L) “LD“– low density, “HD” – high density.
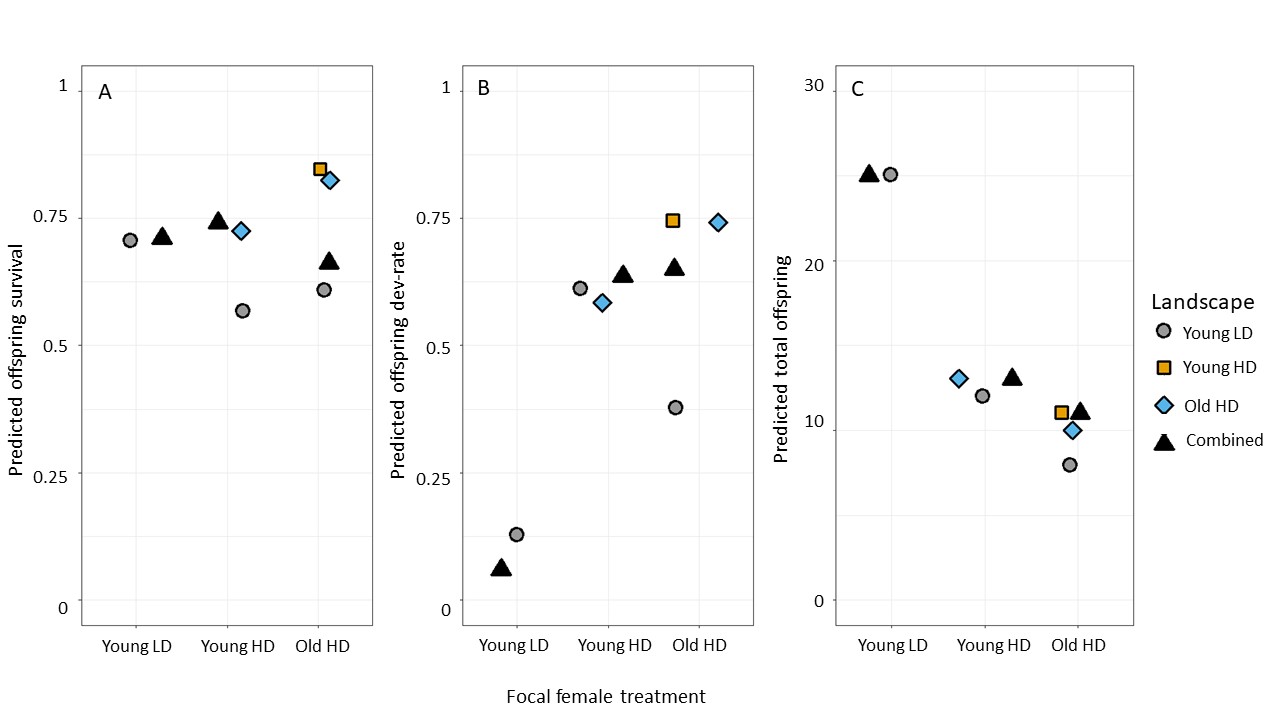

**Figure S10.** Predicted fitness values (y-axis) for (A) offspring survival, (B) offspring development rate and (C) total number of offspring from the individual and combined landscapes shown in Fig S10 for females from young LD, young HD or old HD contexts (x-axis). Predictions made by these four landscapes indicated by different symbols and colours as indicated in the figure legend. If median oviposition behaviour lay outside the bounds of the landscape (see Figure S12), we did not make a prediction.

| **Fitness component** | **Female context** | **Predicted fitness value (from landscape)** | **Observed fitness value** | **Paired t-test, sample estimate** | **p** |
| --- | --- | --- | --- | --- | --- |
| Offspring survival | Young LD | 0.8 | 0.7 | 0.1 | 0.2254 |
|  | Young HD | 0.8 | 0.8 |  |  |
|  | Old HD | 1 | 0.8 |  |  |
| Offspring development rate | Young LD | 0.2 | 0.2 | 0.067 | 0.4266 |
|  | Young HD | 0.5 | 0.3 |  |  |
|  | Old HD | 0.6 | 0.6 |  |  |
| Number of adult offspring | Young LD | 23 | 22 | -1.67 | 0.3701 |
|  | Young HD | 13 | 17 |  |  |
|  | Old HD | 5 | 7 |  |  |

**Table S1.** Summary of paired t-tests for differences between the predicted (from the combined landscape) and observed (experimentally measured) fitness values for young LD, young HD and old HD females.
